# Membrane-bound flurophore-labelled vectors for Targeted DamID allow simultaneous profiling of expression domains and DNA binding

**DOI:** 10.1101/2020.04.17.045948

**Authors:** Caroline Delandre, John PD McMullen, Owen J Marshall

## Abstract

Targeted DamID (TaDa) allows highly efficient cell-type-specific profiling of protein-DNA interactions. Cell-type-specificity, however, is governed by the GAL4/UAS system, which can exhibit differences in expression patterns depending upon the genomic insertion site and the UAS promoter strength. The TaDa system uses a bicistronic transcript to reduce the translation rates of Dam-fusion proteins, presenting the possibility of using the primary ORF within in the transcript to label expression domains and precisely identified the profiled cell populations in experimental samples. Here, we describe new TaDa vectors, pTaDaG, pTaDaG2 and pTaDaM, that use myristoylated-GFP or myristoylated-mCherry as the primary ORF. Differing lengths of the myristoylation sequence between the two GFP plasmids allows additional translational control. The mCherry plasmid allows profiled cells to be visualised when using the NanoDam system, in which an anti-GFP nanobody is fused to Dam to profile the localisation of GFP-fusion proteins. Fly lines created with this system allow easy visualisation of expression domains under both fluorescent dissecting and confocal microscopes without the use of antibody staining, whilst faithfully profiling protein-DNA interactions via Targeted DamID.

## Introduction

Targeted DamID (TaDa) is a recently developed technique that generates *in vivo* cell-type-specific binding profiles of DNA-binding, chromatin-modifying, or DNA-associated proteins in *Drosophila melanogaster* [1, 2]. The technique is highly reproducible and extremely sensitive, generating binding profiles from as few as 10,000 cells in living organisms [2]. The technique is a variant of DamID, in which a protein of interest is fused to DNA Adenine Methylase (Dam) from *Escherichia coli,* leading to the enriched methylation of GATC sites in close proximity to where the protein of interest binds. A recent modification of TaDa, NanoDam, allows the profiling of GFP-fusion protein binding via the fusion of an anti-GFP nanobody to Dam[3].

In TaDa, cell-type-specificity is accomplished via the GAL4/UAS system [4]. High levels of cellular Dam, however, are toxic; and a signature feature of TaDa is a significant reduction in the translation levels of Dam-fusion proteins from highly-expressed GAL4-driven transcripts [1]. In TaDa, this lowering of translation levels is accomplished via a bicistronic transcript, where an upstream ORF is separated from the Dam-fusion ORF by two stop codons and a frameshift. The Dam-fusion ORF is thus only translated via spontaneous ribosome re-initiation, in which the rates of translation of a secondary ORF are inversely proportional to the length of the primary ORF [5]. The original TaDa system used full-length mCherry as the primary ORF. This allows the possibility of visualising the profiled cell population via microscopy of dissected experimental tissue; however, the low brightness of mCherry and the lack of localisation to a particular cellular compartment makes visualisation of expression domains in TaDa experimental samples challenging in practice.

Although broadly defined expression patterns for GAL4 drivers are known, the precise expression pattern and the amount of driver background from other tissues depends upon the targeted insertion site and UAS promoter strength [6]. Given the sensitivity of the TaDa technique and that only a small subset of the cells within an isolated tissue are typically profiled within a TaDa experiment, knowing the exact cells profiled with a GAL4 driver in the TaDa system is critically important to data interpretation.

Here, we describe new TaDa vectors that use membrane-targeted myristoylated-GFP or myristoylated-mCherry as a primary ORF. The pTaDaG vector uses an 85 amino acid (aa) myristoylation sequence; pTaDaG2 and pTaDaM use a 14aa minimal myristoylation sequence. The differences in the primary ORF allow differing secondary ORF translation levels, but otherwise behave identically. The membrane-targeted mCherry plasmid allows visualisation of labelled cells with the recent Nanodam[3] system. Importantly, these primary ORFs allow easy fluorescent identification of profiled cell populations within the experimental sample. The vectors also incorporate a *StuI* restriction site upstream of Dam to easily facilitate the creation of C-terminal Dam-fusion proteins.

## Methods

### Expression constructs

The *pTaDaG* vector was constructed by cutting *pUASTattB* [7] with EcoRI and XbaI, and inserting a 1969bp custom gBlock (IDT) containing *EcoRI-myrGFP-StuI-Dam-MCS-Xbal* via NEB HiFi assembly (NEB). The *myrGFP* sequence represents the first 85aa of *D. melanogaster* Scr64B fused to a *D. melano-gaster*-codon-optimised GFP^F64L,S65T,H231L^ [6]. The *pTaDaG2* vector was created by cutting the *pTaDaG* with EcoRI and NdeI, and inserting a 350bp gBlock (IDT) containing the minimal 14aa myristoylation sequence MGSSKSKPKDPSQR from p60^src^ [8] and the 5’ portion of GFP, again via NEB HiFi assembly. *pTaDaG-Pc* was generated by cutting *pTaDaG* with BglII/XhoI and inserting a 1233bp gBlock (IDT) containing the *Pc-RA* ORF.

*pTaDaM* was generated by cutting *pTaDaG2* with EcoRI / StuI and incorporating an 836bp gBlock (IDT) containing codon-optimised *myr-mCherry,* using the same minimal 14aa myristoylation sequence as TaDaG2.

To create a TaDaM version of NanoDam, *pTaDaM-traNLS-vhhGFP4* was generated by cutting the vector with BglII/XbaI and inserting a 531 gBlock (IDT) containing *traNLS-vhhGFP4,* comprising the nuclear localisation sequence (NLS) from the *tra* gene used in *pStinger* [9] and the vhhGFP4 anti-GFP nanobody sequence[10], codon-optimised for *Drosophila.*

All plasmids were sequence-verified via Sanger sequencing (ABI). Plasmid maps were generated using SnapGene software (Insightful Science).

### Fly lines

GAL4 driver lines used were *worniu-GAL4* [11] for neural stem cells and *R13F02-GAL4* [12] for Mushroom body neurons. Lines were crossed to a *tub-GAL80ts* stock to generate a *worniu-GAL4;tub-GAL80ts* line.

TaDaG-Dam and TaDaG-Pc fly lines were generated by BestGene, Inc (CA), through phiC31-integrase-mediated insertion of the appropriate expression vectors into *attP2* on chromosome 3L.

The TaDaG2-Dam (in *attP2*) and TaDaM-traNLS-vhhGFP (in both *attP2* and *attP40*) were generated via injection of embryos from Bloomington stocks #25709 *(attP40,* Chr2L) and #25710 *(attP2,* Chr3L) that also contained *P{nos-phiC31\int.NLS}* on the X.

### Confocal microscopy

Larval brains (3rd instar, 96hrs ALH) were dissected in PBS and fixed in PBS + 0.3% TritonX-100 (PBST) with 4% (v/v) paraformaldehyde (ProSciTech) for 20 mins, 4°C, before three 10 min washes in PBST. Brains were mounted in Vectorshield + DAPI, and imaged under an Olympus FV3000 confocal microscope at 20x magnification.

### Targeted DamID

TaDaG-Dam or TaDaG-Pc males were crossed to *worniu-GAL4;tub-GAL80ts* virgin females in cages. Embryos were collected on apple juice agar plates with yeast over a 4-hour collection window at 25°C and grown at 18°C for two days. Newly hatched larvae were transferred to food plates for a further five days at 18°C, before shifting to 29°C for 24 hours. Larval brains were dissected in PBS, and processed for DamID-seq as previously described [2, 13] with the following modifications. Briefly, DNA was extracted using a Quick-DNA Miniprep plus kit (Zymo), digested with DpnI (NEB) overnight and cleaned-up with a PCR purification kit (Machery-Nagel), DamID adaptors were ligated, digested with DpnII (NEB) for 2 hours, and amplified via PCR using MyTaq DNA polymerase (Bioline). Following amplification, 2μg DNA was sonicated in a Bioruptor Plus (Diagenode), DamID adaptors removed by AlwI digestion, and 500ng of the resulting fragments end-repaired with a mix of enzymes (T4 DNA ligase (NEB) + Klenow Fragment (NEB) + T4 polynucleotide kinase (NEB)), A-tailed with Klenow 3’ to 5’ exo-(NEB), ligated to Illumina Truseq LT adaptors using Quick Ligase enzyme (NEB) and amplified via PCR with NEBNext Hi-fidelity enzyme (NEB).

The resulting next-generation sequencing libraries were sequenced on a HiSeq2500 and reads were processed with damidseq_pipeline [14].

### Bioinformatic analysis

Binding profiles were visualised using pyGenomeTracks [15]. Heatmaps were generated via the ComplexHeatmap R package [16]. All other plots were generated using R [17].

## Results and Discussion

### Design of the pTaDaG, pTaDaG2 and pTaDaM vectors

All plasmids were generated from *pUASTattB* using synthetic DNA. In designing the *myrGFP* insert for pTaDaG and pTaDaG2, we combined *Drosophila* codon-optimised GFP^F64L,S65T,H231L^ [6] with either the 85aa myristoylation sequence from Scr64B [6] *(pTaDaG)* or a 14 aa minimal p60^src^ myristoylation sequence [8] *(pTaDaG2)* (Fig. 1A). Following the primary ORF, we incorporated a double stop codon / single base frameshift linker, as per the original TaDa vector *(pUAST-attB-mCherry-NDam)* [1], together with a *StuI* restriction enzyme site upstream and in-frame with Dam to allow easy generation of C-terminal Dam fusion proteins (Fig. 1B). Spacing between ORFs is reported to have little effect on translation rates of the secondary ORF [5], allowing the incorporation of the *StuI* site with no translational penalty. The MCS region is separated from Dam using the same Myc-tag+linker sequence as the original vector, allowing cloning compatibility between vectors (Fig. 1C). The pTaDaM vector is identical to pTaDaG2, sharing the same minimal myristoylation sequence, but with codon-optimised mCherry in place of GFP.

**Figure 1:**
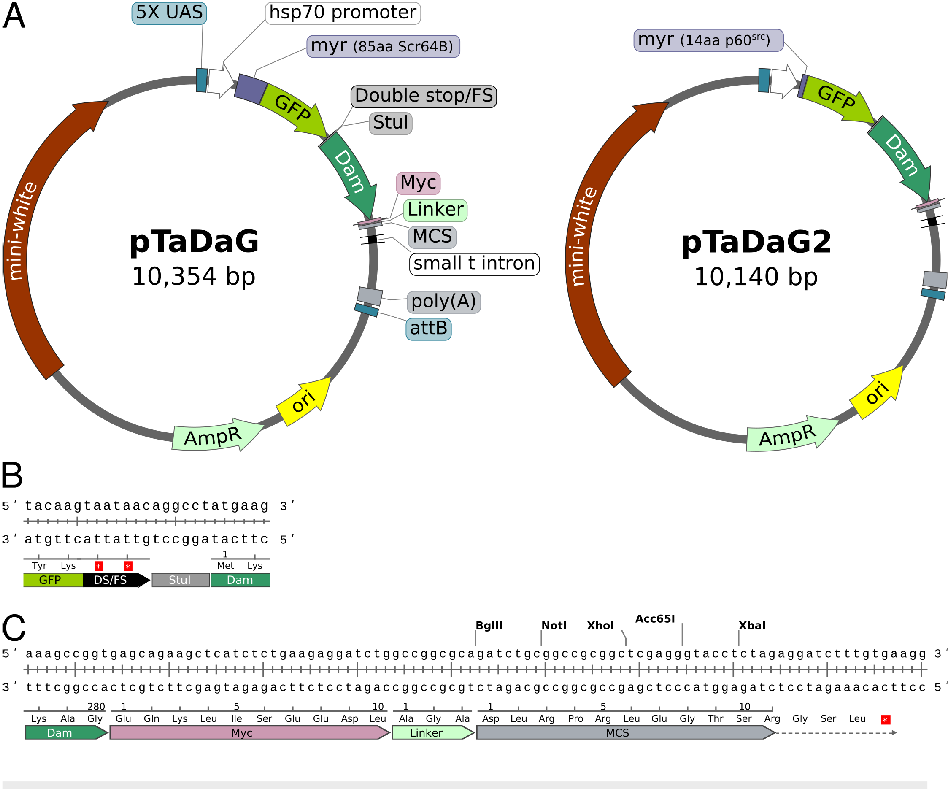
The pTaDaG and pTaDaG2 plasmids. (A) Plasmid maps illustrating the shared features between both plasmids on pTaDaG and the differing myristoylation domains. (B) Sequence of the linker region between the primary GFP and secondary DAM ORFs. (C) Sequence of the Myc-tag-linker-MCS region.

### TaDaG, TaDaG2 and TaDaM label GAL4-driven cell populations *in vivo*

In order to test the labelling capacity of the *myrGFP* primary ORFs in the new vectors, we crossed TaDaG, TaDaG2 and TaDaM-traNLS-vhhGFP flies to the mushroom body neuron driver *R13F02-GAL4* (Fig. 2). All transgenics exhibited clear and specific membranebound fluorescence in mushroom body neurons that was detectable through native fluorescence (without antibody labelling) and was also visible under a fluorescent dissecting microscope (not shown). All variants thus allow simple verification of the expression pattern of Dam-fusion proteins under experimental conditions. The system also allows experimental crosses to be checked for correct cell-type labelling during tissue collection.

**Figure 2:**
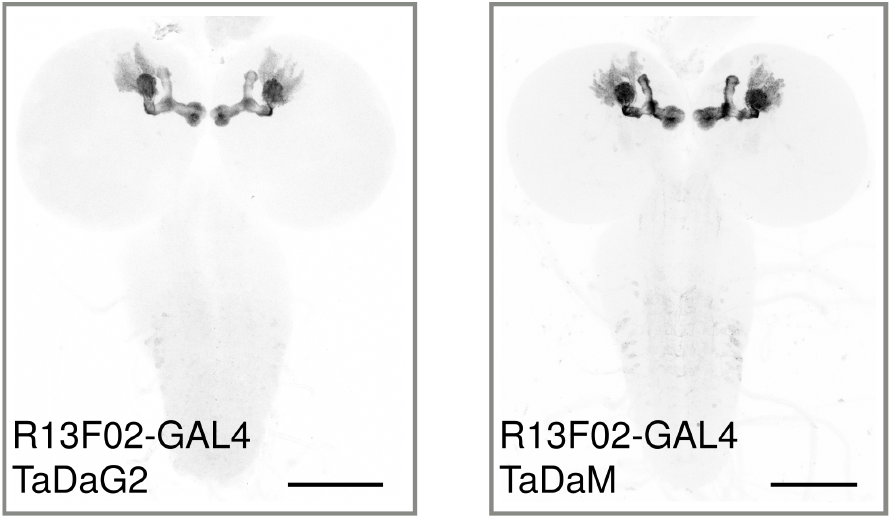
Fluorescent labelling of 3rd instar larval mushroom body neurons via the TaDaG/TaDaM systems. TaDaG2-Dam or TaDaM-traNLS-vhhGFP flies were crossed to the mushroom-body-specific *R13F02-GAL4* driver line and native GFP or mCherry fluorescence imaged. Maximum image projections from confocal stacks are shown. Scale bar: 100μm.

### The TaDaG system faithfully profiles Polycomb binding domains in neural stem cells

To determine if the TaDaG system could generate cell-type-specific Targeted DamID profiles, we profiled Polycomb binding in neural stem cells (NSCs) using the NSC-specific driver *worniu-GAL4,* inducing expression for 24hours in 30 brains (~9000 total profiled neural stem cells) in early (96hrs ALH) 3rd instar larvae (Fig. 3A). Two independent biological replicates had a very high correlation (Pearson’s correlation between TaDaG replicates: 0.92), indicating excellent reproducibility even from very small sample sizes, and no loss of sensitivity when compared to the original TaDa system.

**Figure 3:**
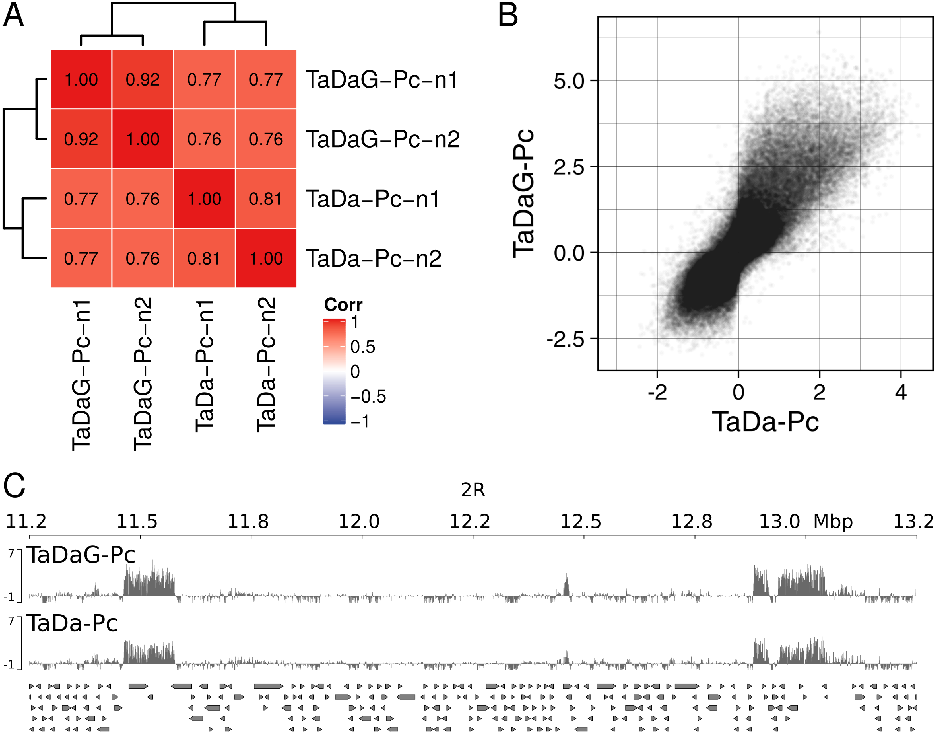
DamID profiling of Polycomb binding in 3rd instar larval NSCs using the TaDaG system. (A) Pearson’s correlation of TaDaG-Pc biological replicates vs previously published TaDa-Pc replicates [13]. (B) Correlation plot of the average Polycomb binding enrichment over all genomic GATC fragments for TaDaG-Pc and TaDa-Pc profiles. (C) TaDaG-Pc and TaDa-Pc binding over two canonical Polycomb foci on chromosome 2R. Scores for (B) and (C) represent normalised log_2_(Dam-fusion/Dam) binding enrichment.

We compared the binding profiles to our previously published Polycomb binding data in NSCs obtained through the original TaDa system [2] (generated using a 16 hour induction timeframe rather than 24 hours in the current study). We observed a high correlation (minimum Pearson’s correlation: 0.77) between the TaDa and TaDaG-generated profiles (Fig. 3A,B) and clear concordant binding over canonical Polycomb foci (Fig. 3C), indicating that the TaDaG system functions indistinguishably from the original TaDa constructs.

Given that GAL4-driver expression patterns are dependent upon both the insertion site and the UAS promoter sequence [6], the ability to verify driver expression in situ is vital for determining the profiled cell population in Targeted DamID and interpreting subsequent binding data. We anticipate that these vectors will prove highly useful to the community.

## Acknowledgments

We thank G. Jefferies for technical assistance. This work was supported by an NHMRC grant APP1128784 and an Ian Potter Foundation equipment grant (20190091) to OJM.

## Notes

### Competing Interest Statement

The authors have declared no competing interest.

### Summary of Updates

A new pTaDaM plasmid is described that uses membrane-bound mCherry as the primary ORF. A pTaDaM-traNLS-vhhGFP variant that can be used for NanoDam is also described. Figure 2 has been revised to show imaging of mCherry in TaDaM-traNLS-vhhGFP fly brains driven with a mushroom-body-specific driver.

